# Monitoring Autophagy: Not All LC3s Are Created Equal

**DOI:** 10.1101/2020.07.20.211458

**Authors:** Yousef A Fouad, Shahenda El-Naggar

## Abstract

The heightened interest in monitoring autophagy comes with discrepant results in different settings. We examined the specificity of reporting in previous studies regarding the isoforms of the most commonly monitored autophagy-related protein, LC3. An ambiguous picture emerged reflecting confusion in design and/or reporting. In light of our current understanding of the different LC3 isoforms, it has become important to specify the targeted isoform when reporting methods and results of studies attempting to monitor autophagy.

## Introduction

Interest in monitoring autophagy in different physiological and pathological settings has significantly increased in the recent years [1]. A simple PubMed search using the term “autophagy” reveals that over 93% of the results exist in the past ten years alone. Despite extensive research, conflicting roles assigned to autophagy within the same setting still exist, prominently in cancer [2]. Part of this conundrum could be attributed to the fact that there is no consensus on the best method to examine autophagy. Each of the available techniques has its own strengths and limitations and only provides part of the full picture [3].

The microtubule-associated protein 1 light chain 3 (MAP1LC3 or LC3 in short, mammalian homologue of yeast Atg8) is a component of the autophagosome and is the most commonly monitored among autophagy-related protiens [3–5]. In mammalian cells, different isoforms exist for LC3 (LC3A, LC3B, and LC3C) with different cellular localization and tissue distribution, and possibly distinct biological functions [3,4]. Accordingly, two problems exist with studies examining LC3 as a marker of autophagy. There is an over-reliance on probing for LC3B in the literature, with no evidence supporting it being the optimal isoform to monitor autophagy [3], and with distinct patterns and contexts emerging for LC3A lately [6–8]. Secondly, there is a lacking (and sometimes confusing) reporting regarding the specificity of the antibodies, constructs, or primers used in the studies against the different isoforms. This could lead to discrepancies between studies and possibly inaccurate conclusions. Here we sample previously published data on the topic to assess such discrepancy and as a cautionary note for new investigators wading into the field of autophagy.

## Methods

We searched the Medline database for studies utilizing LC3 for reporting on autophagy by using the following combination: “light chain 3” OR lc3 OR lc-3 OR “microtubule associated protein 1” OR “microtubule-associated protein 1” OR MAP1 OR MAP-1. The final search was conducted on 7/7/2017 and produced 7437 results. The results were exported into a text file in a numbered latest-to-earliest format. We then randomly selected 10% of the results using a random integer set generator (random.org) producing 744 unique random integers taken from [1, 7437] range. The random integers were matched to their corresponding studies in the exported search results, and full texts were obtained for analysis. We only included primary studies conducted on mammalian cells that examined autophagy using an antibody, construct, and/or primer sequence. Otherwise the study was excluded and a new unique integer corresponding to a new study was randomly generated in place. The analysis of the 744 selected studies included review of both the methods and results sections. The methods section was reviewed for the species of the cells and the reported catalogue number and specificity of LC3-relevant reagents. The results/discussion section was reviewed for the specificity of reporting and its concordance with the respective methods. Reported primer sequences were also reviewed using BLAST (National Center for Biotechnology Information), and the reagents’ specifications were verified using reported catalogue numbers. Data were analyzed using SPSS v23 (IBM).

## Results and Discussion

Of the 744 selected studies, 87.4% (n = 650) were published between 2010 and 2017. Regarding cell species, 59.3% (n = 441) of the studies were conducted on human cells, 21.4% (n = 159) on mouse cells, 17.5% (n = 130) on rat cells, and 1.8% (n = 14) on other mammalian species. Studies that utilized an antibody against LC3 represented 96% (n = 714) of the sample, while 35.1% of the studies (n = 261) utilized an LC3 construct mostly for fluorescence detection of LC3 puncta, and 7.3% (n = 54) utilized LC3 primer sequences for measuring expression levels.

The major trends in the sampled studies utilizing antibodies and/or constructs for detection of LC3 are presented in figure 1. There is a clear underreporting regarding both the catalogue number and specificity of the reagents used. The specific LC3 isoform was reported in less than half of the analyzed studies that utilized LC3 antibodies or constructs (46.4%, n = 331 and 42.5%, n = 111, respectively) creating notable ambiguity. This issue is compounded by possible cross-reactivity between antibodies against different isoforms [4]. Of those reporting the specific isoform in the methods section, only 16.6% (n = 55) for antibodies and 13.5% (n = 15) for constructs followed through with specific isoform reporting in the results. While reporting specific isoforms in methods alone may have previously sufficed, current knowledge about different LC3 isoforms makes it imperative to consistently specify the isoform throughout the manuscript. Noteworthy, four studies mistakenly described detection of a certain LC3 isoform while utilizing an antibody specific to a different isoform.

**Figure 1.**
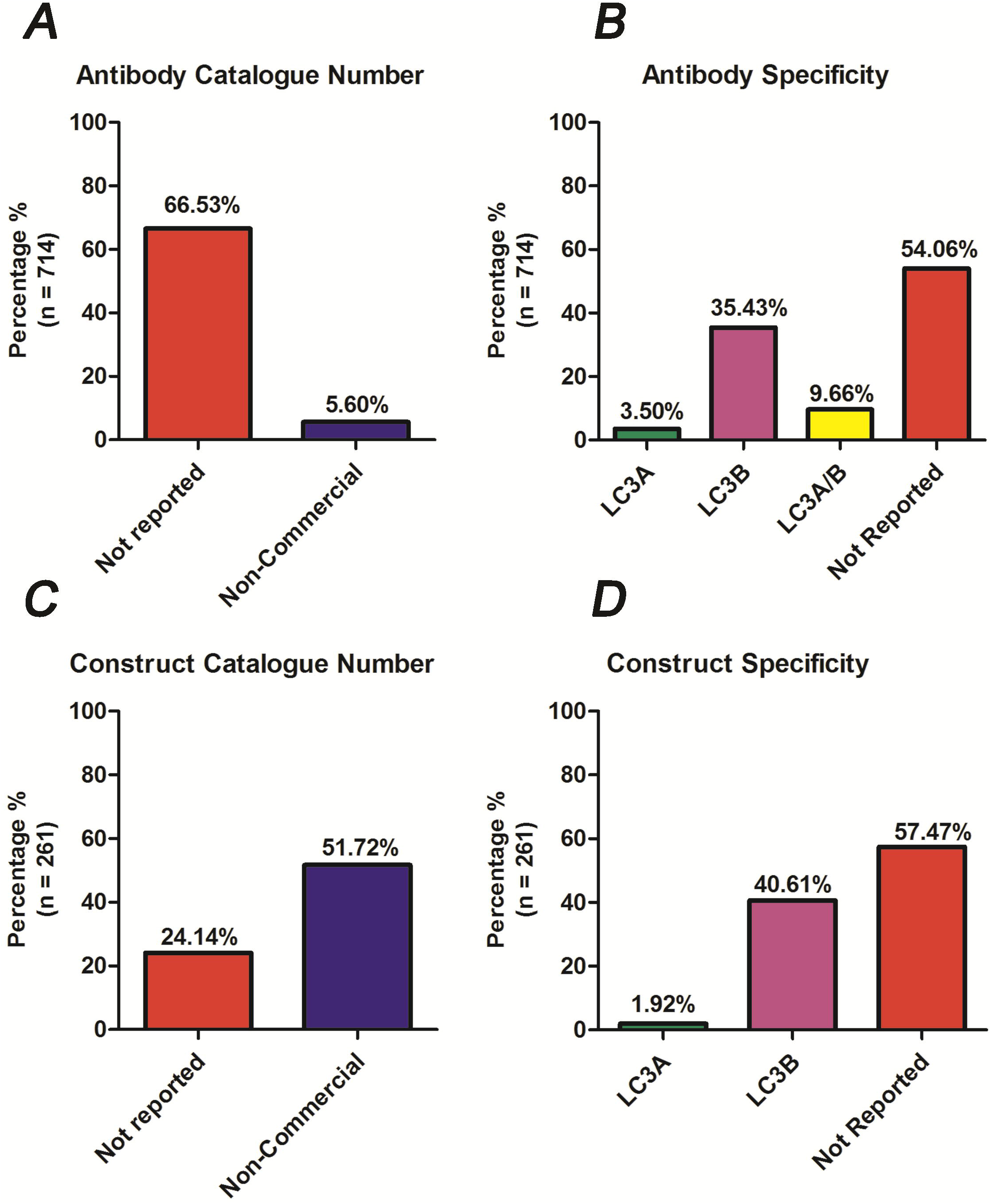
Reporting trends in sampled studies utilizing LC3 antibodies (upper panel) or LC3 constructs (lower panel). A single study may utilize more than one antibody/construct. Non-commercial reagents are those synthesized by either the investigators themselves or another team (obtained as a gift).

As for the 54 studies that adopted a primer design for detection of LC3 expression levels, 18.5% (n = 10) used a sequence specific to LC3A, 74.1% (n = 40) used a sequence specific to LC3B, while 7.4% (n = 4) misused/misreported a sequence that did not correspond to an LC3 region. Yet only 26% (n = 14) described the specific isoform that was targeted while the rest commented on unspecified LC3 levels.

## Conclusion

Autophagy monitoring using LC3 requires being mindful of possible discrepancies arising from the presence of different isoforms. Investigators should accurately report the exact identity and specificity of the LC3 reagent(s) used when describing their methods. They should also be careful not to report total LC3 levels when using an isoform-specific detection technique. Caution should be practiced when dealing with conclusions of studies utilizing LC3 to monitor autophagy and the design should always be thoroughly reviewed.

## Notes

### Competing Interest Statement

The authors have declared no competing interest.

